# Identification of Halo Blight Disease on Oat in Idaho and Exploration of Resistant Sources in Oat, Barley and Wheat

**DOI:** 10.1101/2025.03.05.641278

**Authors:** Jules Butchacas, Suraj Sapkota, Jonathan M. Jacobs, Hannah Toth, Kathy Esvelt Klos, Belayneh A. Yimer

## Abstract

*Pseudomonas coronafaciens* pv. *coronafaciens* (*Pcc*), the causal agent of Bacterial Halo blight (BHB) on oat, has been infrequently reported in the United States, with historical records limited to the 1920s through the 1960s. In 2023, oat trial fields in Aberdeen, Idaho were severely infected with an unknown disease that formed necrotic lesions on leaves. Preliminary identification based on colony morphology suggested a pathogen belonging to the genus *Pseudomonas*. Subsequent whole-genome sequencing confirmed 99.6% average nucleotide identity (ANI) with *Pseudomonas coronafaciens pv. coronafaciens (Pcc)*. This marks the first detection of *Pcc* in Idaho, and the first detailed description of the pathogen in the United States after over half a century. Host range and pathogenicity assessments on multiple cereal crops showed that *Pcc* was pathogenic on oat, barley, and corn. However, wheat, rye and triticale displayed chlorosis and early cell death in response to the pathogen. Evaluation of oat and barley genotypes revealed resistance in the two crop species to be rare with only 2.5, and 4.5% of oat and barley genotypes exhibiting some level of resistance. Notably, the four resistant and moderately resistant barley genotypes identified in this study: DH170472, Celebration, Legacy and Quest are the first to be reported as sources of resistance to BHB. Results of the present study provides a basis for further research toward a better understanding of disease epidemiology, the genetics of host-pathogen interaction and the management of BHB on oat, barley and corn.

## Introduction

Oat (*Avena sativa* L.) is an important multipurpose cereal crop grown worldwide for food, feed, and forage (Ahmed et al. 2014). It is widely acknowledged for its health benefits which includes reducing the risk of cardiovascular diseases (Othman et al. 2011). Furthermore, oat can be grown in marginal lands where other cereal crops do not perform well (Mao et al. 2024). Oat production may be hampered by plant diseases affecting both yield and quality. The common diseases that limit oat production are primarily crown and stem rusts. Other diseases caused by fungi, bacteria, and viruses also impact oats worldwide (Gorash et al. 2017).

Bacterial halo blight (BHB) is an important disease of oats caused by *Pseudomonas coronafaciens* pv. *coronafaciens* (*Pcc*), a Gram-negative bacterium, originally described as *Bacterium coronafaciens,* n. sp. by Elliot (1920) and later reclassified as *Pseudomomas coronafaciens* by Roane (1960). *Pseudomonas coronafaciens* may have also been referred to as *Pseudomonas striafaciens*, *Pseudomonas atropurpurea* and *Pseudomonas zeae* reflecting historical confusion in its taxonomy. The comprehensive revision of *Pseudomonas syringae* taxonomy, presented in the first edition of *Bergey’s Manual of Systematic Bacteriology* (Palleroni, 1984), reclassified 41 nomenspecies as pathovars of *P. syringae*. Among these*, P. coronafaciens*, *P. atropurpurea*, and *P. striafaciens* were reclassified as *P. syringae* pv. *coronafaciens*, *P. syringae* pv. *atropurpurea* and *P. syringae* pv. *striafaciens* respectively. However, the classification of most *P. syringae* pathovars was primarily based on host isolation and restricted cross-pathogenicity testing. Confusion and inaccuracies inherent in phenotype-based taxonomy, particularly in distinguishing closely related subspecies, led researchers to adopt molecular approaches to clarify genetic relationships within the *Pseudomonas syringae* species complex. In a comprehensive DNA-DNA taxonomy study, Gardan et al. (1999) identified nine distinct ‘genomospecies’ within the *P. syringae* species complex. Genomospecies 4 includes seven pathovars, five of which infect domesticated cereals and forage grasses in the Poaceae family. These include *P. syringae* pv. *coronafaciens*, *P. syringae* pv. *atropurpurea*, *P. syringae* pv. *striafaciens*, *P. syringae* pv. *oryzae*, and *P. syringae* pv. *zizaniae*. *P. syringae pv. oryzae has been isolated from rice, while P. syringae pv. zizaniae was isolated from wild rice (Bowden et al 1983; Kuwata. 1985).* Additionally, it includes *P. syringae pv. garcea*, which infects coffee, and *P. syringae* pv. *porri*, which targets onions and leeks. Complementing the genomospecies-based phylogenies that initially rely on DNA-DNA hybridization, multilocus sequence analysis (MLSA) has proven highly effective in resolving the phylogeny of *Pseudomonas syringae* species complex (Berge et al, 2014). This taxonomic approach has allowed the *Pseudomonas syringae* species complex to be divided into 13 distinct phylogroups, with phylogroup 4 being synonymous with genomospecies 4, as it encompasses the seven previously described pathovars. Further whole-genome-based phylogenetic analyses have corroborated the grouping of these seven pathovars within phylogroup 4, reinforcing their close evolutionary relatedness and genetic cohesion (Gomila et al. 2017; Dilon et al. 2019). The species *“P. coronafaciens”* was also proposed by Schaad and Cunfer (1979) based on phenotypic characteristics. Gomila et al. (2017) later provided genomic evidence supporting the reclassification of the seven pathovars within phylogroup 4 as *“P. coronafaciens”*, recognizing it as a distinct nomenspecies. Furthermore, Dutta et al. (2018) combined phenotypic and genotypic analyses, validating the classification of phylogroup 4 pathovars under the species *“P. coronafaciens”*. Collectively, these studies firmly establish *“P. coronafaciens”* as the accepted nomenclature for *P. syringae* phylogroup 4.

The first detailed description of the disease in the United States was published a century ago by Elliot (1920) who documented severe infection of oats throughout Wisconsin during the 1918 season, along with its prevalence in Ohio, Tennessee, California, Virginia, Minnesota, Iowa, Illinois, and Indiana. Although precise data is lacking, BHB can cause substantial economic damage to oats (Elliot 1920; Martens et al. 1984; Roane 1960). In 1960, for example, Roane (1960) reported widespread and severe damage from BHB on oats in Virginia. Since then, limited information is available regarding the disease’s prevalence or confirmed identification of the pathogen in the United States. Elsewhere, the disease has been reported in countries including Brazil, South Africa, Argentina, China, Kenya, South Korea, Japan, Australia, Norway, Sweden, New Zealand and Canada (Chambers and Thomas 2020; Elmhirst 2022; Harder and Harris 1973; Kim et al. 2020a; Malavolta et al. 1997; Persson and Sletten 2021; Smith et al. 1991; Tabei 1964; Tessi 1952; Wang et al. 2024; Wilkie 1972). The widespread geographic distribution of halo blight highlights its adaptability and potential to significantly impact oat production worldwide, emphasizing the need for continued monitoring, comprehensive research, and effective management strategies.

In the summer of 2023, oat trial fields in Aberdeen, Idaho were severely infected by a disease marked by yellowish-brown and necrotic streaks, raising concerns about the emergence of a potential new pathogen (Fig. 1A and B).The emergence of new plant pathogens or strains in regions where they previously did not exit has been increasingly reported worldwide, partly due to intensified global trade (Buja et al. 2021). Therefore, early and accurate diagnosis of such plant pathogens is crucial for effective disease management as the misidentification of a disease and its causal agent can lead to ineffective control measures and wasted resources and time (Riley et al. 2016). The objectives of this study were to 1) identify the causative agent of the new disease observed in the 2023 oat experimental fields in Aberdeen, Idaho, 2) determine the pathogen’s host range, and 3) evaluate potential sources of genetic resistance in selected oat, barley and wheat genotypes to BHB.

**Fig. 1.**
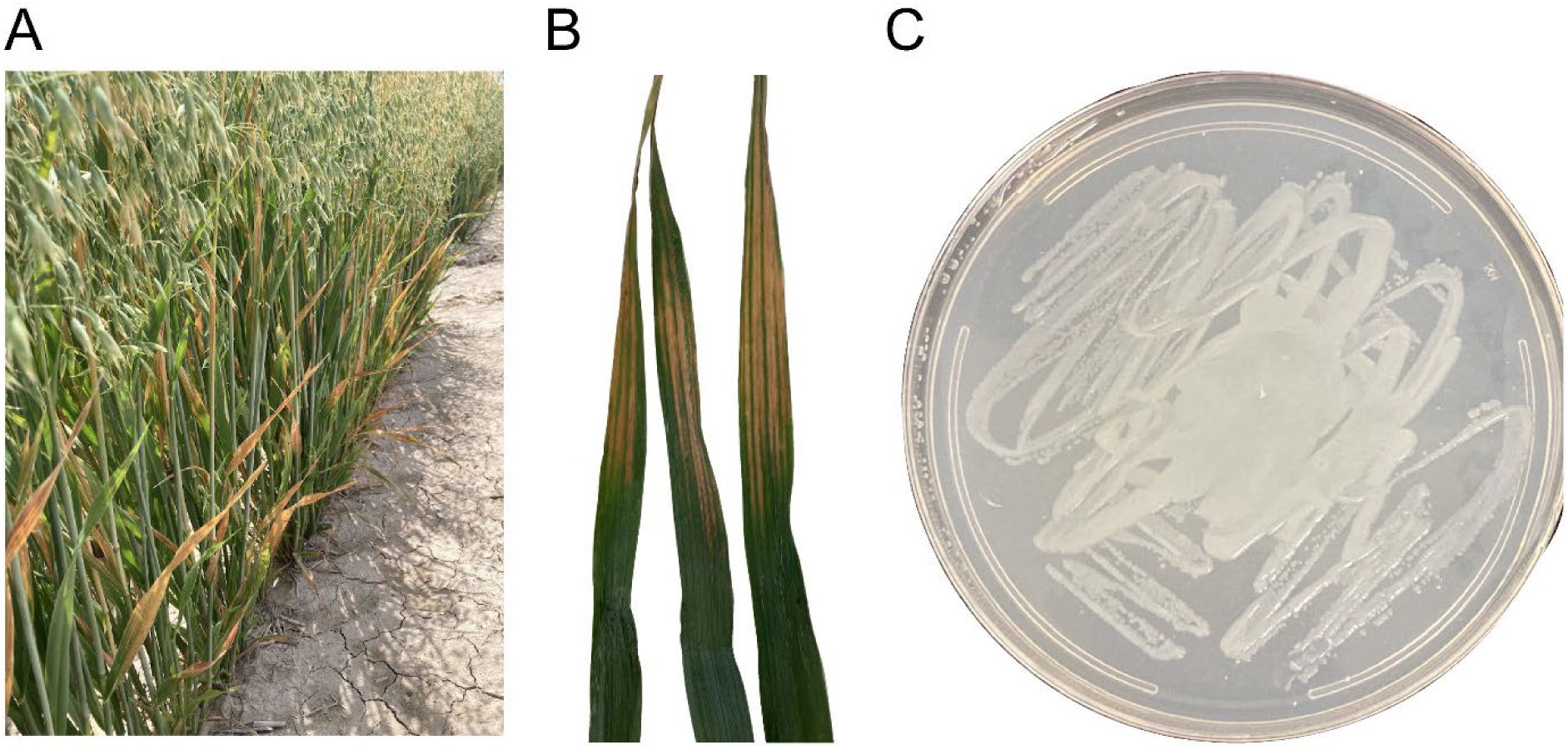
Symptoms of BHB on oat disease caused by *Pseudomonas coronafaciens* pv. c*oronafaciens*. A) in the field in Aberdeen, Idaho in 2023 B) on oat leaves collected from the field C) colonies of *Pcc* on Wilbrink’s media.

## Materials and Methods

### Pathogen isolation

Field-infected oat leaf samples exhibiting brown and oval-shaped lesions were collected and kept in a refrigerator until isolation. To isolate the causal agent of the disease, 1- to 3-cm-long sections of symptomatic leaves from ten different oat plants were surface sterilized with 70% ethanol for 30 s, followed by two rinses in sterile water. The sterilized leaf pieces were then macerated in 300 µl of sterile water on a Petri dish and incubated for 30 min at room temperature. The macerated homogenate was streaked onto Petri dishes containing Wilbrink’s nutrient media (Sands et al. 1986) and kept in an incubator adjusted at 28°C for 48 h. After 48 h, the bacteria were purified by transferring single colonies from each plate onto new plates containing the same nutrient media. The purified bacterial isolates were then used for further analysis.

### Pathogenicity test

The pathogenicity of the bacterial isolate was tested following Koch’s postulates (Bos 1981). Bacterial colonies grown from pure cultures were used to prepare an inoculum suspension with the concentration measured using a DS-C cuvette spectrophotometer (DeNovix Inc., Wilmington, DE), and later adjusted at O.D._600nm_ = 0.4. The oat cv. Idahoat, from which the bacteria were initially isolated was grown in pots in a greenhouse. When plants were at the three-leaf stage, the bacterial suspension was infiltrated into the secondary and tertiary leaves using a 1mL needles syringe. The infiltrated area was marked with a Sharpie for later evaluation. Inoculated plants were kept in a growth chamber adjusted at 26 ± 2^0^C under a 16-hour light period. Five days after inoculation, plants were evaluated for the presence of water-soaked symptoms within the marked area as water-soaking is considered as a compatible reaction to bacterial infection in cereals (Sapkota et al. 2018). Samples from water-soaked lesions were excised and used to re-isolate the pathogen following the procedure described above.

### Molecular diagnosis: Bacterial strains, growth conditions, and DNA extraction

The genome sequences of the 29 *P. coronofaciens* strains used for Average Nucleotide Identity (ANI) analysis were obtained from the National Center for Biotechnology Information (NCBI) GenBank database. The strains represent the 7 different pathovars of the species *Pseudomonas coronafaciens*. These strains were selected based on their infection of a diverse range of hosts, including oat, oatgrass, ryegrass and bromegrass (*pv. coronafaciens,* synonymous with *pv. striafaciens* and *pv. atropurpurea*), domesticated rice (*pv. oryzae)*, wild rice (*pv. zinzaniae)*, coffee (*pv. garcae*), and leek (*pv. porri*). Detailed information for all strains utilized in this study is presented in Figure 2. The 2023 Idaho strain was grown on Wilbrink’s media in an incubator adjusted at 28°C. DNA was extracted from bacterial cultures grown for 48 hours following the QIAGEN Genomic-tip 20/G protocol with the QIAGEN Genomic DNA buffer set. DNA quality and quantity were assessed using a spectrophotometer, measuring 230/280 and 260/280 absorbance ratios.

**Fig 2.**
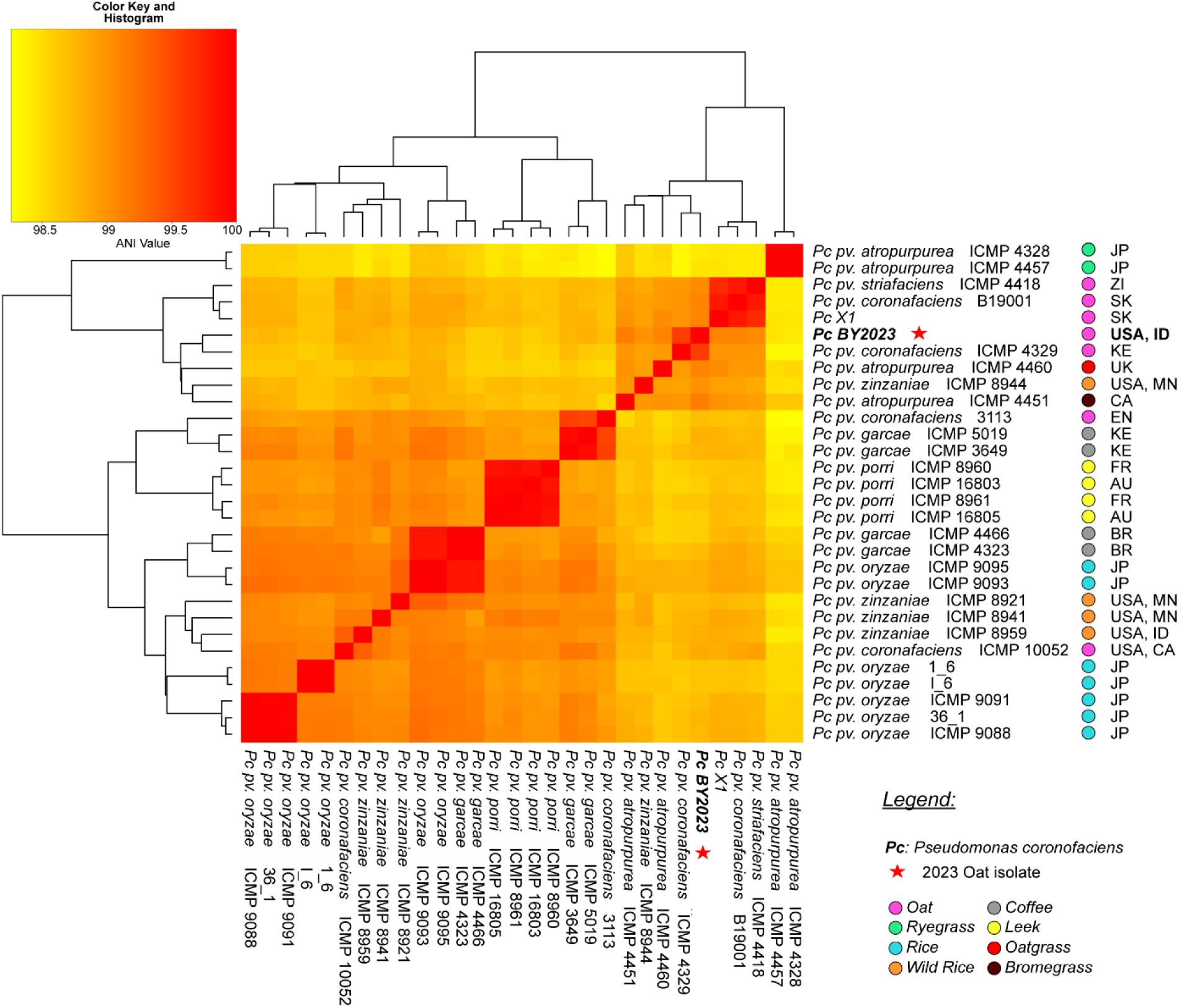
Heatmap and dendrogram of average nucleotide identity illustrating the relationships between ‘*Pcc* BY2023’ and closely related *Pseudomonas coronafaciens* pathovars.

### Molecular diagnosis: Sequencing, genome assembly, annotation and ANI analysis

To accurately identify and classify the causal agent, genomic DNA from the original isolate was extracted and genome sequencing was performed by Plasmidsaurus using long-read sequencing technology provided by Oxford Nanopore Technologies (ONT). The isolate was sequenced twice, and the resulting raw data from both sequencing runs were concatenated. Oxford Nanopore adapter sequences were trimmed using Porechop under the default settings (Wick et al. 2017). Genome assembly was conducted using Flye version 2.9 with parameters set to --genome-size 5m, --iteration 5, and --min-overlap 3000 (Kolmogorov et al. 2019). Polishing was performed using Homopolish, with a mash_threshold of 0.96 (Huang et al. 2021). Genome annotation was conducted with Bakta via the web-based Proksee (Grant et al. 2023; Schwengers et al. 2021). For comparative genomics, an ANI was calculated using PanExplorer (Dereeper et al. 2022) featuring the integrated FastANI module (Jain et al. 2018).

### Host range study

Oat cv. Idahoat, barley cv. Pinnacle, triticale cv. Leaseed, bread wheat cv. Chinese Spring, durum wheat cv. Lure, and a spring rye cultivar were germinated and grown in growth chambers under 16 hours of photoperiod at 22 to 24°C and 70% relative humidity. Corn cv. B73 was germinated and grown under 16 hours of light per day at 28°C and 65% relative humidity. The 1^st^ leaf of 10-day-old seedlings were inoculated as described above with bacterial inoculum of O.D._600nm_ = 0.1. Inoculated plants were kept in a growth chamber at 28°C, under 16 hours photoperiod, and RH of 70%. Reaction of cultivars to *Pcc* infiltration was recorded 2- and 4-days post inoculation (dpi).

### Evaluation of oat, barley and wheat genotypes for resistance to BHB

A total of 161 oat (129 breeding lines, 21 landraces and 11 varieties), 89 barley (55 varieties, 31 breeding lines and 3 landraces) and 41 wheat (35 breeding lines and 6 varieties) genotypes were evaluated for their reaction to *Pcc* isolate BY2023 under controlled conditions at the seedling stage. Seeds were planted in cone-tainers of 3.8 mm diameter × 210 mm depth (Stuewe and Sons, Inc., Tangent, OR) filled with a 3:2:2 mix of sand, peat moss, and vermiculite.

The Osmocote plus 15-9-12 fertilizer (ICL, St. Louis, MO) was applied to the soil mix before planting. Each line was planted in two cones, two seeds per cone, and the experiment was repeated once. After planting, the cones were placed in a greenhouse at 23-26°C with a 16-hour photoperiod. Preparation of the bacterial culture and inoculation was done as described by Sapkota et al. (2018) with some modifications. The pure isolate of *Pcc* stored in a −80 °C freezer was streaked into Wilbrink’s agar (WBA) plate and grown at 28 °C for 48 h. The bacterial cells were collected from the WBA plate using a sterilized loop and suspended in sterile water. The inoculum concentration was measured using a DS-C spectrophotometer and adjusted to an O.D._600nm_ = 0.4. A needles syringe was used to inject the bacterial suspension into the middle section of the fully developed secondary leaf. Two infiltration spots were made in each leaf and the area where the inoculum was infiltrated was marked using a Sharpie marker. After inoculation, plants were kept in a growth chamber with a temperature of 26 ±2°C and a photoperiod of 16 hours. Disease rating was done 5- days after inoculation using a scale based on the amount of water-soak developed. The rating scale was defined as follows: Resistant (R) = necrosis of the marked area with no water-soaking; Moderately Resistant (MR) = necrotic area with low level of water-soaking on the periphery of the infiltrated area; Moderately Susceptible (MS) = water-soaking limited within the infiltrated area; Susceptible (S) = water-soaking of the infiltrated area and the symptom extended beyond the infiltrated area (Fig. 4).

## Results

### Morphological and pathogenic characterization of the causative agent

Isolation of the causal agent from infected oat leaves resulted in the growth of typical creamy-white bacterial colonies on WBA, a morphological characteristic of *Pseudomonas* spp. (Fig. 1C). Koch’s postulates confirmed the causal relationship between the isolated creamy-white bacterial colonies and the observed disease symptoms. Inoculation of healthy oat plants with the purified bacteria reproduced the characteristic water-soaked lesions seen in the field. Re-isolation from these lesions yielded colonies identical in morphology and growth characteristics to the original isolates, conclusively identifying the bacterium as the disease-causing agent.

### Taxonomic identification of the causative agent

Whole-genome sequencing (WGS) using Oxford Nanopore Technologies (ONT) was employed for precise taxonomic identification of the 2023 oat isolate, referred to as ‘*Pcc* BY2023’, hereafter. Average Nucleotide Identity (ANI) analysis was conducted, comparing the isolate to 29 representative strains from the seven pathovars of the *P. coronafaciens* species. The genome of ‘*Pcc* BY2023’ revealed a high degree of similarity with *Pseudomonas coronafaciens* pv. *coronofaciens* and its synonymous taxa *pvs. striafaciens* and *atropurpurea*, with ANI values exceeding 99% (Fig. 2). Notably, *Pcc* BY2023 demonstrated the closest genetic relationship to the *pv. coronafaciens* oat isolate ICMP 4329 from Kenya, with an ANI of 99.57%. Based on these results, BY2023 can be taxonomically designated as *Pcc*. Sequence data was deposited under NCBI Bioproject ID: PRJNA1178904; accession JBISUX000000000.

### Pathogenicity and host range

The pathogenicity and host range assays were conducted on a panel of cereal crops: oat, barley, rye, triticale, durum wheat, bread wheat, and corn. The results indicated that *Pcc* BY2023 was pathogenic on oat, barley, and corn. Symptoms on oat and barley appeared at 2 dpi, manifesting the water-soaked lesions. By 4 dpi, these water-soaked lesions on both oat and barley evolved into sunken, gray-brown tissue, indicating a progression towards more severe necrosis. In contrast, corn did not exhibit water-soaked lesions at 2 dpi instead, it developed grayish, sunken lesions from the onset, which became more extensive and coalesced by 4 dpi. Rye, durum wheat, and bread wheat initially displayed chlorosis and early cell death at 2 dpi, which transitioned to a hypersensitive response (HR) characterized by a chlorotic halo surrounding the dead tissue by 4 dpi. Triticale showed a mild chlorotic response at 2 dpi, which became more pronounced by 4 dpi (Fig. 3).

**Fig. 3.**
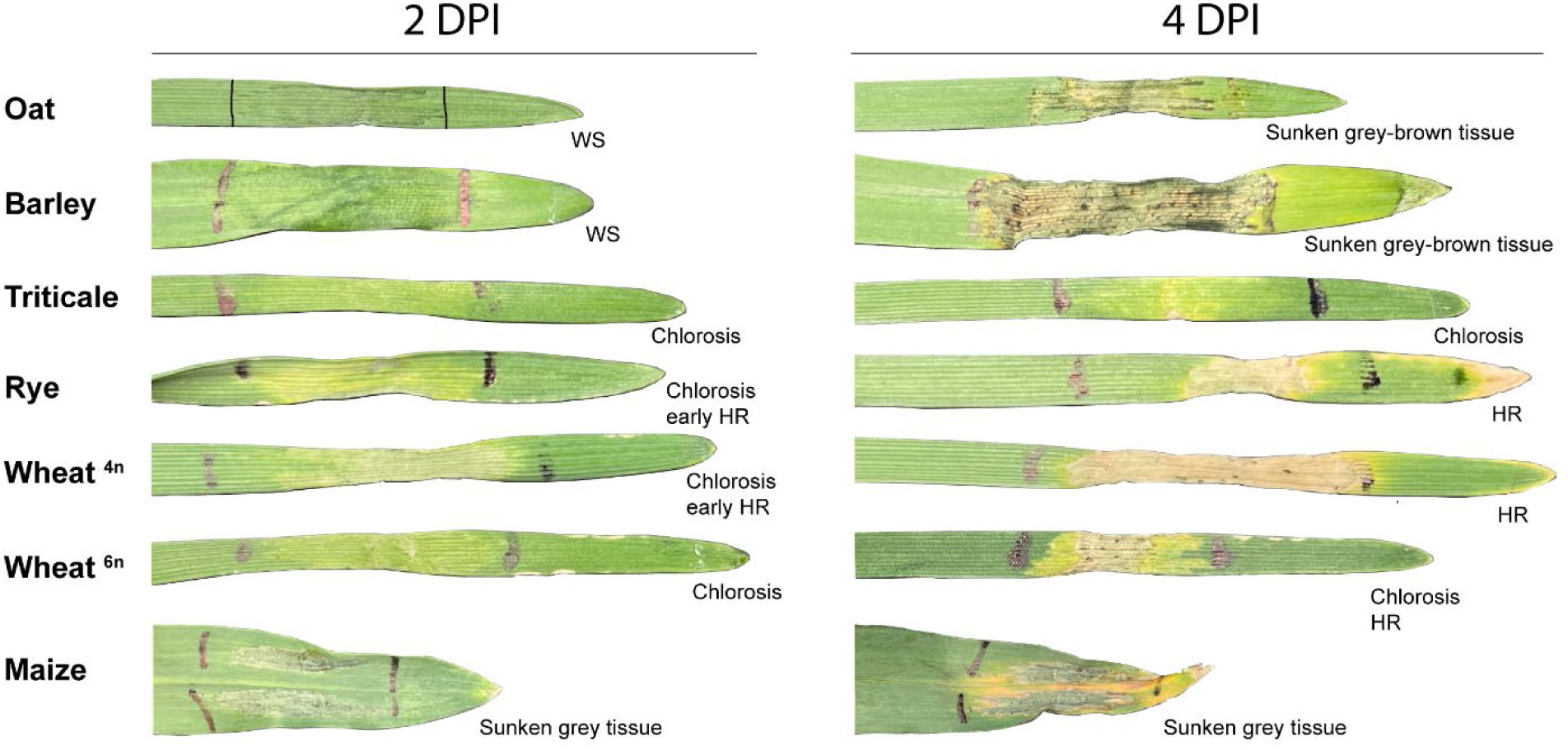
Host response of cereal crop species to infection by *Pseudomonas coronafaciens* pv. *coronafaciens* under controlled environment. Oat, barley, triticale, rye, wheat (4n and 6n) and corn plants inoculated at the first leaf stage with O.D._600nm_=0.1 concentration of bacteria and evaluated two and four days post inoculation. WS = Water-Soaking; HR = Hypersensitive Response

**Fig. 4.**
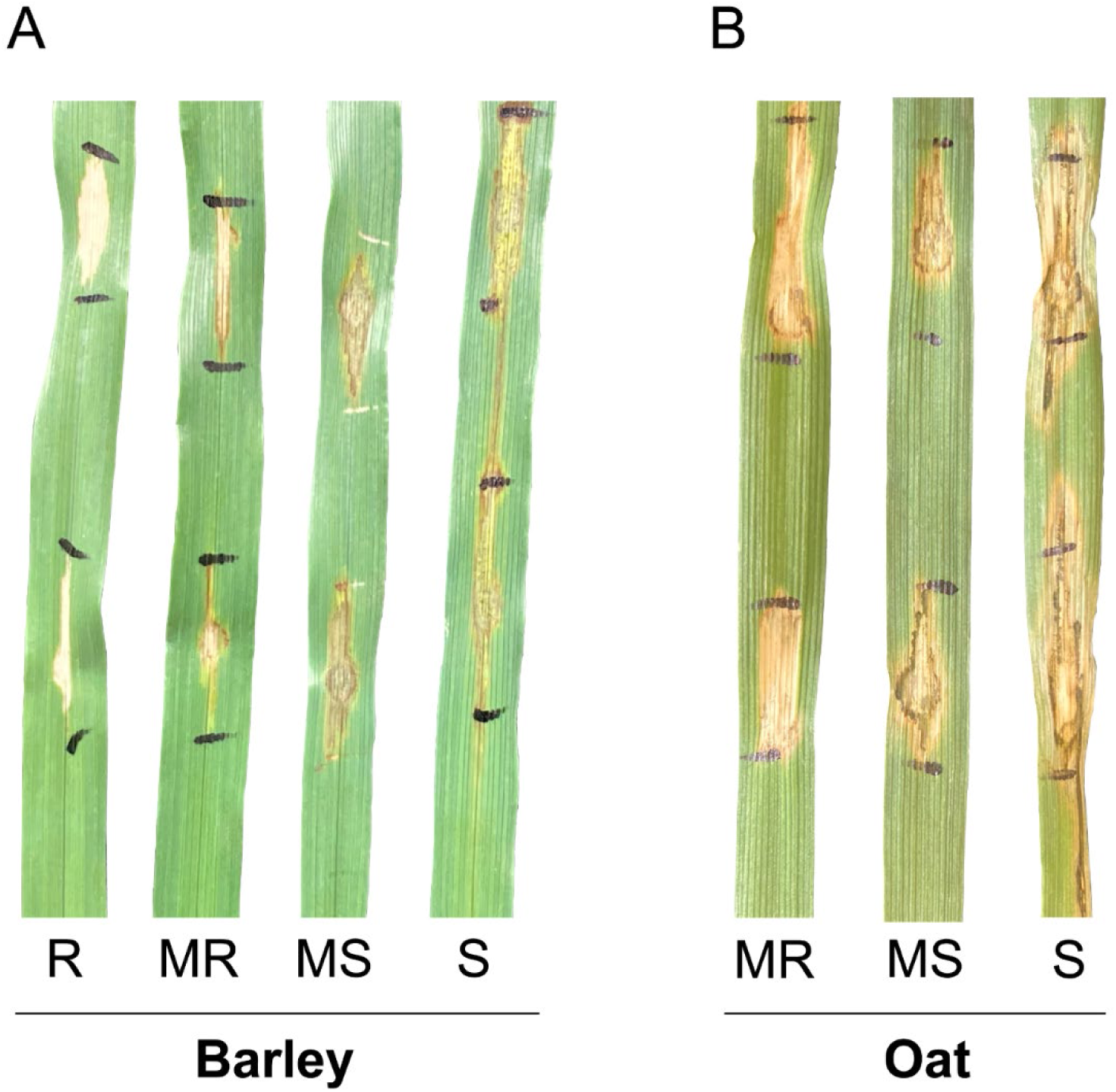
Disease scale used to evaluate oat, barley, and wheat genotypes against *Pseudomonas coronafaciens* pv. *coronafaciens*. A) symptoms on barley, B) symptoms on oat: Resistant (R) = necrosis of the marked area with no water-soaking; Moderately Resistant (MR) = necrotic area with low level of water-soaking on the periphery of the nfiltrated area; Moderately Susceptible (MS) = water-soaking limited within the infiltrated area; Susceptible (S) = water-soaking of the infiltrated area and the symptom extended beyond the infiltrated area.

### Reaction of oat, barley and wheat genotypes to BHB

None of the oat genotypes evaluated in this study were resistant. Four genotypes (2.5%): ‘FL22618’, ‘IN_CHILE_22-03’, ‘UCD-5’, and ‘TAM-O-401’ developed a relatively low level of water-soaking demonstrating moderately resistant reaction in both experiments. Ninety-six oat genotypes (59.6%) were moderately susceptible. On 61oat genotypes (37.9%), the water-soaking symptom extended beyond the infiltrated area indicating a susceptible reaction. In barley, a doubled haploid line (DH170472) from Oregon State University had a resistant reaction in both trials. Three other genotypes: ‘Celebration’, ‘Legacy’ and ‘Quest’ had moderately resistant reaction. Most of the barley genotypes (93.3%) had moderately susceptible reaction. Only two barley genotypes exhibited the water soaking symptom beyond the infiltrated area indicating a susceptible reaction to the disease. The reaction of wheat genotypes to *Pcc* BY2023 was consistent with the initial pathogenicity test. All wheat genotypes exhibited either resistant (75.6%) or moderately resistant (24.4%) reactions (Table 1). Reaction of all oats, barley and wheat genotypes is presented in Table S1 A - C.

**Table 1.** Infection response of oat, barley and wheat to *Pseudomonas coronafaciens* pv. *coronafaciens*.

| Crop species | Genotypes<br>evaluated (#) | Infection response |  |  |  | Some resistance<br>(%) |
| --- | --- | --- | --- | --- | --- | --- |
|  |  | R | MR | MS | S |  |
| Oat | 161 | 0 | 4 | 96 | 61 | 2.5 |
| Barley | 89 | 1 | 3 | 84 | 2 | 4.5 |
| Wheat | 41 | 31 | 10 | 0 | 0 | 100 |

R = resistant MR = moderately resistant MS = moderately susceptible S = susceptible

## Discussion

The first description of BHB on oats was presented by Galloway and Sotjhworth (1890). However, the causative pathogen was first isolated and described by Elliot (1920). The last detailed report of the disease in the United States was decades ago in Virginia (Roane 1960). Since then, there has been no detailed description or report about its prevalence or damage in the United States. This study combined traditional methods, including colony morphology and pathogenicity tests, along with advanced genomic tools to accurately diagnose the causative agent. Furthermore, selected genotypes were evaluated for resistance, providing insights into potential use of host resistance to manage BHB. The present study marks the first documented occurrence of BHB on oat in Idaho, and the first detailed description of the pathogen in the United States since 1960. Further research is needed to fully understand the factors contributing to the appearance of the disease in Idaho. However, it could be potentially linked to the increased use of pivot irrigation, which might have created a favorable microenvironment for disease infection and development. This possibility was also suggested as contributor to the appearance of Fusarium head blight on barley and wheat in Idaho (Bissonnette et al. 2018) and to the increased incidence and severity of bacterial leaf streak (BLS) on barley (Marshall et al. 2024; *Belayneh Yimer*, personal observation). The prevalence of BHB on oat in Idaho warrants a thorough scouting and survey in major oat-producing regions of the United States, mainly in the Midwest, where the environmental conditions are more favorable for *Pcc* infection and disease development.

Whole genome sequence has emerged as an essential tool for accurate taxonomic classification and identification of plant pathogens (Tambong et al. 2022). In addition to precise pathogen identification, WGS enables the detection of pathogenicity and virulence factors, making it invaluable for plant pathogen research (Sapkota et al. 2020). The ANI analysis results provide compelling evidence supporting the taxonomic identification of the 2023 oat isolate as *Pcc*, a pathovar traditionally designated for oat-infecting strains. *Pcc* BY2023 demonstrated the closest genetic relationship to the oat isolate *Pcc* ICMP 4329 from Kenya, with an ANI of 99.57%. Seven other strains shared ANI values exceeding 99% with *Pcc* BY2023, including three pv. *coronafaciens* strains (originating from oat), one pv. *striafaciens* strain (originating from oat), two pv. *atropurpurea* strains (originating from oatgrass and bromegrass), and one pv. *zizaniae* strain (originating from wild rice). Richter and Rosello-Mora (2009) recommended the use of an ANI boundary of >95% for taxonomically circumscribing prokaryotic species. These findings suggest that the host range-based pathovar system may not be well-suited for the taxonomy of this species as it encompasses genetically similar strains with overlapping and, in some cases, distinct host ranges. Host range, the defining criterion for pathovar designation, is influenced by plant-pathogen compatibility, which can be determined by the presence or absence of single effector genes (e.g., avr genes). This system fails to account for the genetic continuity and evolutionary relatedness within strains that exhibit minor variations in host specificity, leading to artificial distinctions between pathovars. Therefore, strains designated as different pathovars within the *P. coronafaciens* species share high genetic similarity but are classified separately based on their primary host plants. Conversely, some strains within the same pathovar group fall into distinct genetic clusters, underscoring the inconsistencies and limitations of host range-based classifications in accurately reflecting genetic relationships within this species. This discrepancy highlights the necessity of adopting a genomic framework, to complement or replace traditional pathovar designations, enabling a more accurate and biologically meaningful taxonomy. Implementing genome-based taxonomy for the *Pseudomonas coronafaciens* species would offer a more robust and consistent classification system, better reflecting the close genetic relationships among strains while accounting for the plasticity of the host range driven by minor genetic variations. Such an approach would not only enhance the resolution of evolutionary and taxonomic studies but also support functional research and disease management efforts by focusing on genetic determinants, including those linked to pathogenicity and potential host-specific resistance traits.

The presented study highlighted a complicated host-pathogen interaction than previously supposed. Elliot (1920) observed that under natural environment only oat was infected by BHB. However, when inoculation was done in a controlled environment condition, barley, rye, and wheat were infected but not corn. Similarly, a study conducted in China to determine the host range of *Pcc* found that out of six different plant species, including corn, only oat was susceptible to infection by *Pcc* (Wang et al. 2024). The present study indicated that *Pcc* is pathogenic to oat and barley in a controlled environment concurring with previous studies. However, unlike the two previous studies, *Pcc* was pathogenic on corn (Fig. 3). This aligns with the findings of Shaad and Cunfer (1979), who demonstrated that pv. *coronafaciens* (including the former pv. *zeae*), pv. *atropurpurea*, and pv. *striafaciens* could infect a broad range of cultivated cereals, including corn. The observed discrepancies in corn pathogenicity among *Pcc* strains may be attributed to variations in specific effector genes involved in virulence or avirulence, which play a pivotal role in determining host compatibility. Alternatively, these differences could result from the use of distinct corn genotypes in various studies, with some genotypes exhibiting resistance while others are susceptible to *Pcc*. This highlights the necessity of further research to identify the molecular mechanisms underlying the resistance and susceptibility of corn to *Pcc*. Such investigations are crucial for understanding the pathogen’s epidemiological potential, especially given the importance of corn as a major crop in the United States. Addressing this knowledge gap is essential for preparedness in managing potential outbreaks and safeguarding corn production.

Another interesting observation was the interaction between *Pcc* and barley. In the location where BHB was observed in 2023, there were barley plots adjacent to oat plots. These barley plots were infected by BLS caused by *Xanthomonas translucens* pv. *transclucens*, but not by BHB However, in the controlled environment study, the same barley varieties that were planted adjacent to oat plots were susceptible to BHB. In an earlier study by Elliot (1920), oat and barley were planted so close together that the plants were intermingled at the margins. Interestingly, only the oat plants were heavily infected, and no disease was observed on the barley. The author concluded that while *Pcc* may occasionally infect wheat, rye and barley slightly, it rarely, if ever, appeared on anything but oats in the field. It warrants further investigation to assess whether this conclusion remains valid, as shifts in pathogen virulence and potential host jumps may have occurred over time. Nevertheless, host range studies have consistently shown that barley is highly susceptible to *Pcc* under controlled conditions, while for corn, rye, and wheat, the interactions vary, with both compatible and incompatible responses observed depending on the study. Although natural infections of barley by *Pcc* have never been observed, the consistent susceptibility of barley to the pathogen in artificial inoculations remains unexplained. Artificial inoculations, such as syringe infiltrations, bypass natural plant defenses by injecting high bacterial concentration directly into the mesophyll. In contrast, natural infections involve low bacterial numbers entering through natural entry points, requiring the pathogen to be highly adapted to both the host and its specific environment to establish a significant population and cause disease. *Pcc* may be highly adapted to naturally infect oats but not the other alternative cereal hosts. Further investigation into the disease ecology and the pathogen’s fitness on potential alternative hosts, especially in barley, corn, and rye, is needed.

Host plant resistance is the most preferred method to manage plant diseases. In small grain cereals, evaluations of germplasm for resistance to bacterial diseases is done using wounding or non-wounding methods. Infiltration of bacterial suspension using needleless syringes is commonly practiced under controlled conditions (Curland et al. 2018; Sapkota et al. 2018; Tambong et al. 2024). However, under field conditions, it is common to directly spray high concentration bacterial inoculum on to plant leaves (Kandel et al. 2012; Ritzinger et al. 2023). In this study, we adopted the infiltration method utilized by various authors (Kim 2020b; Sapkota et al. 2018; Wang et al. 2024) to evaluate the reaction of oat, barley and wheat genotypes to *Pcc*.

In oat, no genotypes were immune or highly resistant to BHB. This may be due to the small number of genotypes evaluated. However, this is not unique to oats. For example, wheat, barley and triticale germplasm highly resistant to bacterial leaf streak are rare (perhaps 1 – 2%) (Kandel et al. 2012; Ritzinger et al. 2023; Sapkota et al. 2018). In the present study only four oat genotypes (2.5%) had a moderate level of resistance, which suggests that BHB resistance in oats may be rare. Additional efforts to evaluate germplasm for resistance could yield additional sources of moderate resistance and may identify stronger resistance than observed in this study. In the meantime, the moderately resistant oat genotypes identified in the present study can serve as starting materials to develop BHB resistant oat germplasm and to map the underlying genetic loci.

Evaluation of oat germplasm during the 1940s and 1960s identified several varieties with resistance to BHB, i. e., ‘Victoria’, ‘Bond’, ‘Fulghum’ (Kingsolver 1944), ‘Benton’, ‘Cherokee’, ‘Clinton’ (Caldwell et al. 1946), ‘Cc4146’ (Griffiths and Peregrine 1964), ‘Victorgrain’ and ‘Dubois’ (Cheng and Roane 1968). Most of these varieties were used extensively as parents in oat improvement programs near the time of their published characterization. Within the oat genotypes evaluated for this study were nine lines from the National Small Grains Collection annotated as descending either from ‘Victorgrain’, ‘Dubois’ or ‘Cc4146’. All except one were susceptible in both trials of this study while ‘CIav4832’ was moderately resistant in the first test but susceptible in the next (Table 2). If these accessions were developed without selection for halo blight resistance, the loss of such is not unexpected. Alternately, the lack of resistance may be due to a shift in pathogen virulence arising over time or across geographic regions as the earlier studies were conducted half a century ago.

**Table 2.** Previous and current reaction of some oat genotypes to BHB caused by *Pseudomonas coronafaciens* pv. *coronafaciens*.

| Accession | Name | Previous reaction | Current reaction |  |
| --- | --- | --- | --- | --- |
|  |  |  | Test 1 | Test 2 |
| Ciav 5353 | Victorgrain | R<sup>a</sup> | MS | MS |
| CIav 5354 | Victorgrain | R<sup>a</sup> | MS | MS |
| Ciav 5114 | Victorgrain | R<sup>a</sup> | MS | MS |
| Ciav 4832 | Victorgrain | R<sup>a</sup> | MR | MS |
| Ciav 4685 | Victorgrain | R<sup>a</sup> | MS | MS |
| Ciav 6572 | Dubois | R<sup>b</sup> | S | MS |
| PI 592075 | Cc4146 | R<sup>c</sup> | S | MS |
| PI 592076 | Cc4146 | R<sup>c</sup> | MS | MS |
| PI 410850 | Cc4146 | R<sup>c</sup> | MS | MS |

R = resistant MR = moderately resistant MS = moderately susceptible S = susceptible
<sup>a</sup> = Kingsolver 1944 <sup>b</sup> = Cheng and Roane 1968 <sup>c</sup> = Griffiths and Peregrine 1964

Most importantly, to the best of our knowledge, this is the first study to evaluate barley germplasms and identify resistance to BHB in barley. Like oat, the genetic resistance in barley seems rare as only 4.5% of the genotypes were either resistant or moderately resistant. Since BHB has never been observed to infect barley under field conditions, it may not be a priority disease for further research. However, the resistance sources reported here could be utilized to develop resistant germplasm if the disease becomes important on barley. The reaction of wheat to *Pcc* infection needs further investigation to conclusively determine whether the observed reaction is a resistant or a nonhost reaction.

In summary, the foliar disease symptoms of oat plants observed at the USDA-ARS experimental research plots in Aberdeen, Idaho in 2023 was caused by *P. coronafaciens* pv. *coronafaciens*, the causal agent of bacterial halo blight. To the best of our knowledge, this is the first report of BHB of oat caused by *Pcc* in Idaho, USA. Additionally, the host range of *Pcc* was determined using a panel of cereal crops, including oat, barley, rye, triticale, durum wheat, bread wheat, and corn. Evaluation of a set of oat, barley and wheat genotypes for resistance to BHB identified four moderately resistant oat genotypes, and one and three resistant and moderately resistant barley genotypes, respectively. Results of the present study may provide a basis for further research toward a better understanding of the disease epidemiology, the genetics of host-pathogen interaction, and overall management of BHB.

## Acknowledgements

This research was partly supported by the U.S. Department of Agriculture, Agricultural Research Service. Mention of trade names or commercial products in this publication is solely for the purpose of providing specific information and does not imply recommendation or endorsement by the U.S. Department of Agriculture. USDA is an equal opportunity provider and employer.

